# Early life growth and cellular heterogeneity in the short-lived African turquoise killifish telencephalon

**DOI:** 10.1101/2024.05.01.591995

**Authors:** Caroline Zandecki, Valerie Mariën, Rajagopal Ayana, Jolien Van houcke, Lutgarde Arckens, Eve Seuntjens

## Abstract

The African turquoise killifish (*Nothobranchius furzeri*) is becoming a favorable model for neurobiological research. The combination of a short lifespan and a declining neuroregenerative capacity upon aging makes the killifish ideally suited for research on brain aging and regeneration. A remarkable cellular diversity makes up the young-adult killifish telencephalon, characterized by highly proliferative non-glial progenitors and four spatially distinct radial glia subtypes. In contrast to a relatively slow embryonic development, hatching is followed by a period of accelerated growth in the short-lived *N. furzeri* strain; GRZ. Accordingly, the brain of this teleost experiences a period of rapid expansion and maturation. In this study, we charted the growth progression of the killifish telencephalon during early post-embryonic development. We identified a dynamic cellular buildup of the neurogenic niches sustaining this explosive growth. Spatial data revealed specific differences between pallial and subpallial regions in terms of growth pace and cellular output. Spatial signatures comparable to zebrafish were identified for excitatory and inhibitory neuronal lineages, already present at hatching.

**Summary statement:** This study charts the progenitor and neural diversity of the telencephalon from post-embryonic development, identifying the cellular building blocks during accelerated brain growth of the short-lived African Turquoise killifish.

## Introduction

The African Turquoise killifish (*Nothobranchius furzeri)* has developed into an established model for age-related research, in part because of its remarkably short natural lifespan. Annual *Nothobranchius* fishes inhabit temporary water ponds in Southeast Africa. The unpredictable and temporally restricted alternations of rainy and dry seasons pressured these fishes to develop explosive growth and attain fast sexual maturation (Valdesalici and Cellerino, 2003; Blažek *et al*., 2013; Reichard and Polačik, 2019). The GRZ *N. furzeri* strain has the shortest lifespan recorded and maintains this characteristic under laboratory conditions (Valdesalici and Cellerino, 2003; Cellerino *et al*., 2016; Van houcke *et al*., 2021).

Research on killifish has been carried out to study the impact of aging on various biological processes and systems such as the immune system (Bradshaw *et al*., 2022), circadian rhythm (Barth *et al*., 2021), epigenetics (Zupkovitz *et al*., 2021), wound healing (Örling *et al*., 2023) and the central nervous system (Tozzini *et al*., 2012; Van houcke *et al*., 2021; Vanhunsel *et al*., 2021). This revealed that, in contrast to other highly regenerative species, the brain of this short-lived teleost loses its impressive regeneration capacity upon aging, and even adopts mammalian traits, such as glial scar formation upon injury (Wendler *et al*., 2015; Van houcke *et al*., 2021; Vanhunsel *et al*., 2022). These human-like aging features of the killifish central nervous system have prompted researchers to investigate the occurrence of spontaneous neurodegeneration and other age-related disease hallmarks for diseases like Parkinson’s disease and Amyotrophic Lateral Sclerosis (Valenzano *et al*., 2006; Matsui *et al*., 2019; Bagnoli *et al*., 2022; Louka *et al*., 2022; Bergmans *et al*., 2023; de Bakker and Valenzano, 2023).

The period of explosive growth after hatching is preceded by a very slow embryonic development in the killifish. Depending on the time spent in diapause, embryonic development can take 3-4 weeks to multiple years (Genade *et al*., 2005; Blažek *et al*., 2013; Api *et al*., 2018). This is in stark contrast with the extremely fast development of common teleost model systems such as zebrafish (*Danio rerio*), goldfish (*Carassius auratus*) and medaka (*Oryzias latipes*), with an embryonic development period of 2,3 and 9 days, respectively (Kimmel *et al*., 1995; Iwamatsu, 2004; Tsai *et al*., 2013). After fast embryonic development, zebrafish mature slowly into a sexually mature adult in approximately three months and do not experience a period of accelerated growth (Parichy *et al*., 2009). Killifish, on the contrary, reach sexual maturity in 5-6 weeks. The growth of the killifish pallial surface, for example, is reported to be three times faster than that of the zebrafish in the first two weeks of development (Coolen *et al*., 2020), but the growth pattern and appearance of progenitors and neural cell types from hatching to adult have not been documented yet.

The neurogenic niches and neural progenitor cells are identified for the adult killifish brain and comparisons could be made with established model systems like zebrafish and mouse. In the telencephalon, three neurogenic regions are identified, as was done for zebrafish (Adolf *et al*., 2006; Grandel *et al*., 2006; Tozzini *et al*., 2012). Region I originates in the subpallium and is homologous to the mammalian subventricular zone. Two pallial regions are described to be homologous to the mammalian hippocampal subgranular zone, with region II stretching over the pallial surface and region III located at the posterior zone of the dorsal pallium (Tozzini *et al*., 2012). In zebrafish, radial glia (RG) are the main progenitor cell type and are responsible for the production of neurons both during development and in the adult brain (Furlan *et al*., 2017; Coolen *et al*., 2020), although heterogeneity in the progenitor population is also reported in zebrafish (Grandel *et al*., 2006; März *et al*., 2010). The progenitors responsible for the bulk of proliferation in the killifish telencephalon, termed non-glial progenitors (NGPs), lack the typical RG transcriptional profile yet possess an apical domain and basal process (Coolen *et al*., 2020; Ayana *et al*., 2023). NGPs are present already in the young post-hatching killifish brain and can be recognized by expression of PCNA, MSH1, HMGB2A and STMN1A (Coolen *et al*., 2020; Ayana *et al*., 2023). Our previous single-cell transcriptomics and spatial analysis further revealed a tremendous progenitor cell diversity in the adult killifish telencephalon. Next to two NGP subtypes, we identified four RG subtypes and multiple intermediate cell states. The RG subclusters display an astroglial (Astro-RG1/Astro-RG2, marked by SLC1A2 and/or CX43 respectively), neuroepithelial (NE-RG3, marked by ZIC2) or ependymal (EPD-RG4, marked by EPD) transcriptional profile and all have distinct spatial locations in the adult telencephalon (Ayana, Zandecki *et al*., 2023).

Since previous killifish research has mainly focused on aging, unique adaptations necessary for accelerated growth in the early life stages may have been overlooked. In this study, we investigated the growth progression of the killifish telencephalon during early post-embryonic development and identified the pattern of growth and cellular heterogeneity present during this explosive growth phase. We additionally reflected on the differences and similarities with zebrafish and other relevant model systems.

## Materials and Methods

### Animals and housing

All experiments were performed with African Turquoise killifish (*N. furzeri*) from the inbred GRZ-AD strain. When the embryos were ready to hatch (Golden Eye stage), the eggs were moved to a hatching tank with a small volume of ice-cold hatching solution (Humic acid, Sigma-Aldrich, 53680, 1g/L in aquarium water) with a continuous flow of oxygen (Polačik *et al*., 2016; Van houcke *et al*., 2021). The fish were raised in the hatching tank for one week at 26°C with a daily addition of fresh aquarium water. After the first week, juvenile fish were transferred to a 3.5 L aquarium in a zebTEC Multi-Linking Housing System (Techniplast) and housed under standardized conditions: temperature 28°C, pH 7, conductivity 600 μs, 12h/12h light/dark cycle. After hatching, the fish were fed twice a day with *Artemia salina* (Ocean Nutrition). From four weeks onwards, they were fed with *Artemia salina* (Ocean Nutrition) and mosquito larvae (*Chironomidae*). All experiments were approved by the KU Leuven ethical committee (025/2021; 155/2023), under the European Communities Council Directive of 20 October 2010 (2010/63/EU).

### EdU labelling

To label dividing cells and trace the progeny, batches of fish (n=3-6) were placed in 5-ethynyl-2’-deoxyuridine (EdU, 4 mM) for 16h. Each group was pulsed once with EdU at a specific age: 1-day post-hatching (dph), 1 week (w), 2w, 3w, 4w and 5w. All fish were sacrificed at the age of 6w (Fig. 2A).

### Tissue collection and processing

Hatchlings (1-5dph), juveniles (2w) and young-adult fish (6w) were euthanized in 0.1% buffered tricaine (MS-222, Sigma Aldrich). Hatchling and juvenile tissue (full body) were fixed overnight in 4% paraformaldehyde (PFA, 8.18715, Sigma-Aldrich, in phosphate-buffered saline (PBS)) at 4°C. After euthanasia, young-adult fish were perfused with PBS and 4% PFA (Mariën *et al*., 2022). The brains were dissected and fixed overnight in 4% PFA at 4°C. Next, the tissue (full body or brain) was saturated overnight in 30% sucrose (in PBS) after which the tissue was embedded in 30% sucrose and 1.25% agarose in PBS. 10 µm-thick coronal sections were cut with a CM3050s cryostat (Leica) and the sections were collected on SuperFrost Plus Adhesion slides (10149870, Thermo Fisher Scientific). The sections were stored at - 20°C until HCR, IHC or EdU staining.

### Hybridization chain reaction (HCR)

The probe pools used for HCR were generated and validated as described in detail before (Deryckere *et al*., 2021; Van houcke *et al*., 2021; Elagoz *et al*., 2022; Styfhals *et al*., 2022) and ordered via Integrated DNA Technologies, Inc (IDT). The HCR protocol (HCR v3.0), based on the protocol of Choi *et al*., (2018), is adapted for cryosections as described in Van houcke *et al* (2021), without the Proteinase K permeabilization step. To optimize the signal-to-noise ratio, the probe pool concentration ranged between 0.3-1.8 pmol, depending on the target (Suppl. Table 1). In case the amplifier (Molecular Instruments, Inc) was linked to Alexa Fluor 546, the sections were mounted with SlowFade Gold Antifade Mountant (S36936, Invitrogen) to improve the photostability of the fluorophore. When combining HCR with IHC, the slides were washed 3 times with 5xSSCT (0.1% Tween-20 in saline-sodium citrate buffer) after the hybridization and amplification steps, before proceeding with the IHC protocol.

### Immunohistochemistry (IHC)

The cryosections were first dried for 30 minutes at 37°C before proceeding with wash steps in TBS (0,1% Triton-X-100). The tissue was blocked with 20% normal donkey serum (D9663, Sigma-Aldrich) in TNB (Tris-NaCl blocking buffer). After blocking, the sections were incubated overnight with the primary antibody diluted in Pierce Immunostain Enhancer (46644, Thermo Fisher Scientific) (Suppl. Table 2). After wash steps in TBS, the sections were incubated with the secondary antibody in TNB for 2 hours. Afterwards, the sections were rinsed in PBS followed by a nuclear staining with DAPI (4’,6-diamidino-2-fenylindool, 1:1000 in PBS, 32670, Sigma-Aldrich). All sections were mounted with Mowiol and a glass cover slide. Whenever IHC was preceded by an HCR, all wash steps were performed with PBS-T (0.1% Tween-20 in PBS) instead of TBS.

### EdU staining

The EdU staining was performed using the Click-iT™ EdU Cell Proliferation Kit for Imaging (C10340, Invitrogen). Briefly, the sections were dried for 30 minutes, rinsed in PBS and permeabilized in 0.5% Triton X-100 in PBS. After an additional wash step in 3% BSA (bovine serum albumin in PBS), the Click-iT® reaction cocktail was incubated for 30 minutes in the dark. Afterwards, sections were rinsed with 3% BSA, and a nuclear stain with DAPI was performed. In case the EdU staining was combined with an IHC, the EdU staining preceded the IHC. The sections were mounted with Mowiol and a glass cover slide.

### Imaging

Images were acquired with a widefield (Axio Observer7, ZEISS) or confocal (LSM 900 with Airyscan 2, ZEISS) microscope. The images were further processed with ZEN 3.7 software (ZEISS). The distances (D) of the telencephalon (Suppl. Fig. 2A) were measured using the Line tool in ZEN 3.7.

### Section selection

The selection of sections of comparable anterior-posterior levels of the different stages (1dph, 5dph, 2w, 6w) was based on anatomical landmarks and the distance between sections (in µm). The approximate distance between the sections in Fig. 1 and Suppl. Fig. 1 are the following: (1dph) 50µm, 50µm; (5dph) 50 µm, 100µm; (2w) 100 µm, 100 µm; (6w) 180 µm, 120 µm, from rostral to caudal. The following anatomical landmarks were used: the olfactory bulb is visible in rostral sections; middle sections were selected based on the progenitor region extending from the dorsal to ventral surface and the presence of a specific ventrolateral neuronal cluster; in caudal sections, a central circular neuronal cluster is visible in the pallium, comparable to the central zone of the dorsal pallium as described in the adult killifish telencephalon (D’angelo, 2013).

**Figure 1:**
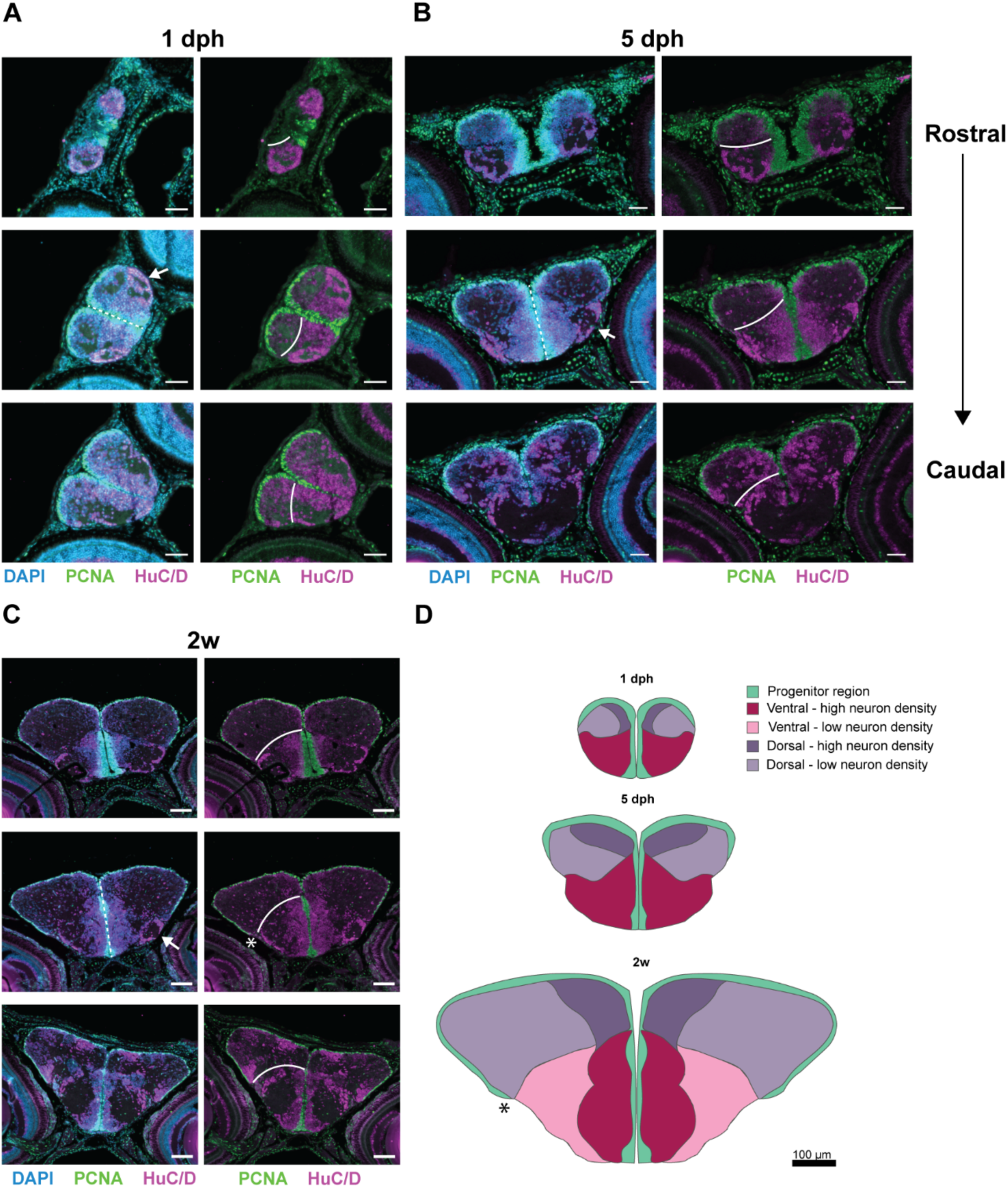
Explosive growth of dorsal and ventral telencephalic domains. Immunohistochemical staining for PCNA (green) and HuC/D (magenta) in combination with a nuclear stain (DAPI, blue) on coronal brain sections. Three levels of the anterior-posterior axis are shown for **(A)** 1dph, **(B)** 5dph, and **(C)** 2w old killifish, presented from rostral to caudal. White line represents the pallial-subpallial border. **(D)** On scale illustration summarizing the distribution of progenitor (green) and neuronal regions (shades of red/purple) on coronal telencephalic sections of 1dph, 5dph, and 2w-old juveniles. Example sections of a comparable anterior-posterior level were selected based on the presence of a specific ventrolateral neuronal cluster (arrow (A-C) and the progenitor region extending from the dorsal to the ventral surface (dashed line (A-C)). **(A-B)** Scale bar: 50 µm, **(C-D)** Scale bar: 100µm. **Abbreviations:** dph: days post-hatching, w: week.s

## Results

### Explosive growth in the post-embryonic telencephalon

To chart the source of post-hatching growth and morphogenesis of the telencephalon, we first visualized the proliferative progenitor and postmitotic neuronal domains at different stages from hatching to young-adult age (6w). We used the proliferation marker PCNA to visualize cell division at 1 dph, 5 dph, 2w and 6w, in combination with the pan-neuronal marker HuC/D (Fig. 1A-C, Suppl. Fig. 1A-B). At 1dph and 5dph, PCNA^+^ progenitors occupied one continuous region across the pallial surface from medial to lateral (Fig. 1A-B). The olfactory bulb was not directly covered with a progenitor layer at the dorsal side, visible in the rostral-most section at 1dph (Fig. 1A). This aligns with what is known in the developing zebrafish, where proliferating cells cover the dorsal surface between 2-5 days post fertilization (dpf), but stay absent dorsally of the olfactory bulb (Folgueira *et al*., 2012). At 2w, we detected that the PCNA^+^ pallial progenitor layer expanded further into the posterior pallial region (Fig. 1C-D, asterisk). The major anatomical structures of the adult telencephalon could clearly be recognized at 2w, yet the telencephalon still undergoes a massive expansion from 2w onwards. This could be observed when comparing the juvenile (2w) telencephalon to the young-adult (6w) telencephalon (Suppl. Fig. 1A).

To visualize the absolute difference in overall size of the telencephalon upon development, we created a representative illustration at the mid-anterior-posterior level of all ages discussed (Fig. 1D, Suppl. Fig. 1B). We further delineated the growth progression from hatchling to young-adult by measuring the horizontal and vertical expansion of the pallium and subpallium (Suppl. Fig. 2). These absolute numbers indicate a greater expansion of the pallium compared to the subpallium in all directions measured. In the first week post-hatching, the dorsal telencephalon (Fig. 1D, purple, pallium), started expanding laterally and grew beyond the lateral subpallial edge (Fig. 1D, pink region). The progenitor region covering the dorsal surface stretched along the expanding pallium. Stretching of the pallial surface in the first two weeks post-hatching was already quantified by M. Coolen and colleagues (2020) and revealed that the dorsal surface increased 5-fold in this period (Coolen *et al*., 2020). From 1dph to 2w, we measured a 4-fold increase on average in the pallium, confirming their results (Suppl. Fig. 2). The subpallium, on the other hand, was proportionally bigger than the pallium at hatchling stages (1-5dph) and was very dense in neurons compared to the pallium (Fig. 1A-B, the pallial-subpallial border is indicated with a white line). The growth of the subpallium decreased between 2w and 6w compared with juvenile stages (1dph-2w) and is paralleled by an increase in neuropil and extracellular matrix (Suppl. Fig. 1). The surface of the proliferative zone of the subpallium did not stretch out as much as the pallium while the telencephalon increased in size. In general, we concluded that the pallial growth (10-fold increase) contributed most to the massive expansion of the telencephalon (overall 7-fold increase) from hatching to young-adult.

### Birthdating reveals two different stacking processes within the telencephalon

To understand whether there was a temporal order in the generation of neurons and whether they could be found in organized patterns throughout the telencephalon, we performed an EdU birthdating experiment (Fig. 2A). The progeny of the dividing progenitors at the time of EdU pulse (1dph, 1w, 2w, 3w, 4w and 5w) was always visualized at 6w of age (Fig. 2B). Cells born early after hatching (from 1dph) were found in the parenchyme in a concentric circle parallel to the pallial surface, leaving the center of the 6w pallial parenchyme devoid of EdU^+^ cells. Labeling at later ages revealed neurons in consecutive circles at closer distances to the ventricular surface (Fig. 2B). This indicates that the center region of the pallium was already formed during embryonic development, similar to the dorsal pallium in zebrafish (Furlan *et al*., 2017). Subpallial growth occurred in a comparable consecutive manner. A dense line of EdU^+^ cells parallel to the subpallial neurogenic region could be identified, with the distance to the midline inversely related to the time of EdU tracing (Fig. 2B). We also observed a scattered pattern of EdU^+^ cells throughout the subpallium, indicative of tangential migration in the subpallium.

**Figure 2:**
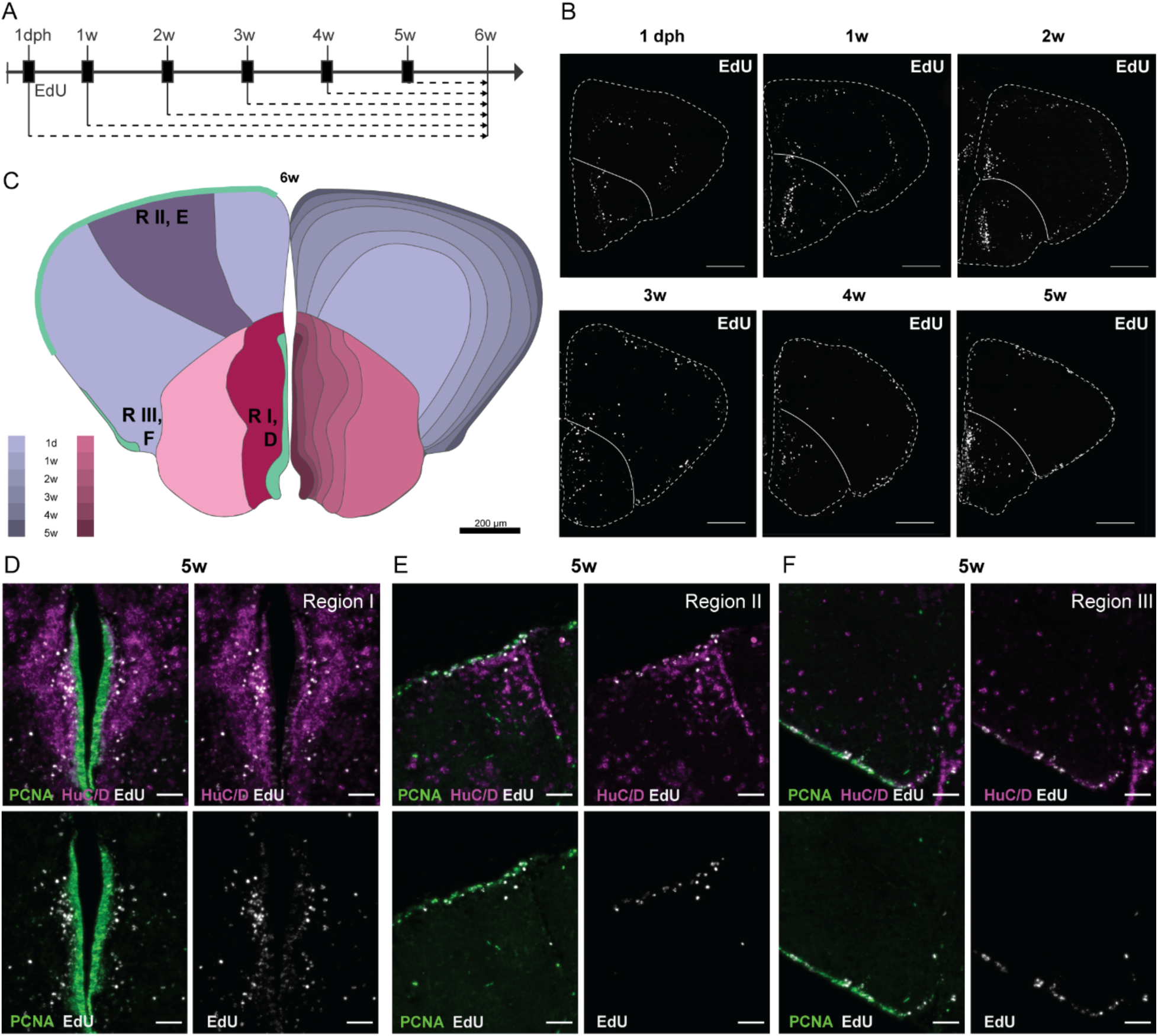
EdU tracing reveals a neuronal stacking process during telencephalic growth. **(A)** Design of the EdU birthdating experiment. Each experimental group is exposed to 4mM EdU for 16h (black box) at different time points during development. EdU is incorporated into the DNA of dividing cells, their progeny is always traced until 6w. **(B)** Distribution of EdU^+^ nuclei (white) on coronal sections at the mid-anterior-posterior level at 6w. The timepoint of EdU administration is indicated above each panel. The outer border of the sections is indicated with a dashed line and the pallial-subpallial border with a full line. In the pallium, EdU^+^ cells appear in concentric circles at differing distances from and parallel to the pallial surface. In the subpallium, the EdU^+^ cells appear more scattered, but the bulk of EdU^+^ cells seem to have migrated a comparable distance from the subpallial ventricular zone for each timepoint of EdU treatment. Scale bar: 200 µm. **(C)** Illustrative summary of the EdU tracing experiment. The opposite stacking process of the pallium and subpallium is color-coded for each period. The location of the pictures in panels D-F (neurogenic region I, II, III) is indicated on the left hemisphere. **(D-F)** Zoom-in on EdU^+^ cells at each neurogenic region upon EdU treatment at 5w. The EdU staining (white) is combined with an immunohistochemical staining for HuC/D (magenta) and PCNA (green). After 1 week of tracing from 5w, the EdU signal is visible in the PCNA^+^ progenitor cells and HuC/D^+^ neurons at a distance from the ventricular zone. Note that Region I, contains more EdU^+^/HUC/D^+^ cells than Region II and III. A varying degree of EdU intensity is observed in the PCNA^+^ progenitors.

Systematic analysis of multiple coronal sections at the mid-anterior-posterior axis for each experimental group is summarized on top of the 6w old telencephalic representation (Fig. 2C). Each layer demonstrates the estimated marginal growth during this period for the pallium and subpallium separately. Neurogenesis seems to be additive from hatching and persists similarly into adulthood. From the thickness of the layers (Fig. 2C) and the distance of EdU^+^ cells from the ventricular surface (Fig. 2B), we can infer that the number of cells generated at later time points decreased.

In general, subpallial neurogenic region I is known to have the highest level of proliferation compared to pallial regions II and III (Tozzini *et al*., 2012; Ayana *et al*., 2023). At 5 weeks, strongly labeled EdU cells that were born at the time of labeling, were observed at a distance from the progenitor zone in region I, while in region II these cells were still found in the progenitor zone, suggesting that in the subpallium, postmitotic cells move away from the neurogenic niche faster than in the pallium (Fig. 2D-F). A high level of PCNA^+^ progenitor cells were EdU^+^ in the subpallium, albeit in lower intensity, indicative of their faster proliferative rates (asymmetric divisions) during the one-week tracing period. In the pallium, a lower number of HuC/D^+^/EdU^+^ neurons were present, in line with a lower proliferating capacity of neurogenic region II and III (Fig. 2E-F). Next to the PCNA^+^ progenitors with low intensity EdU, we could observe PCNA^+^ progenitors with high intensity EdU signal, which could indicate that these progenitors did not divide further during the tracing period of one week, or certainly at a slower rate compared to the progenitors in the subpallium.

The killifish telencephalon grows by addition of cells at the dorsal ventricular surface and the ventral midline. However, this stacking process is not the only factor contributing to telencephalon growth. From hatching to young-adult age (6w), a large increase in the formation of neuropil was observed for both pallial and subpallial domains, at the expense of nuclear density (Fig. 1A-C, Suppl. Fig. 1A). Hence, it is the combination of proliferation, migration and maturation that mitigates the explosive growth of the developing telencephalon.

### Differential organization of radial glia cells in the developing telencephalon

A radial glia scaffold is essential for the proper neuroanatomical development of the telencephalon as RG fiber structures provide a scaffold for axon growth and neural migration to their final location in the whole network (Rakic, 1971; Kaur *et al*., 2020). At 5dph, RG-cell bodies and fibers were visible upon SLC1A2 (GLT1/EAAT2) mRNA detection (Fig. 3A-C). In the pallium, a radial trajectory of the RG fiber structures was visible with the RG cell bodies located in the ventricular zone. A comparable radial fiber structure was present at 2w (Suppl. Fig. 3A).

**Figure 3:**
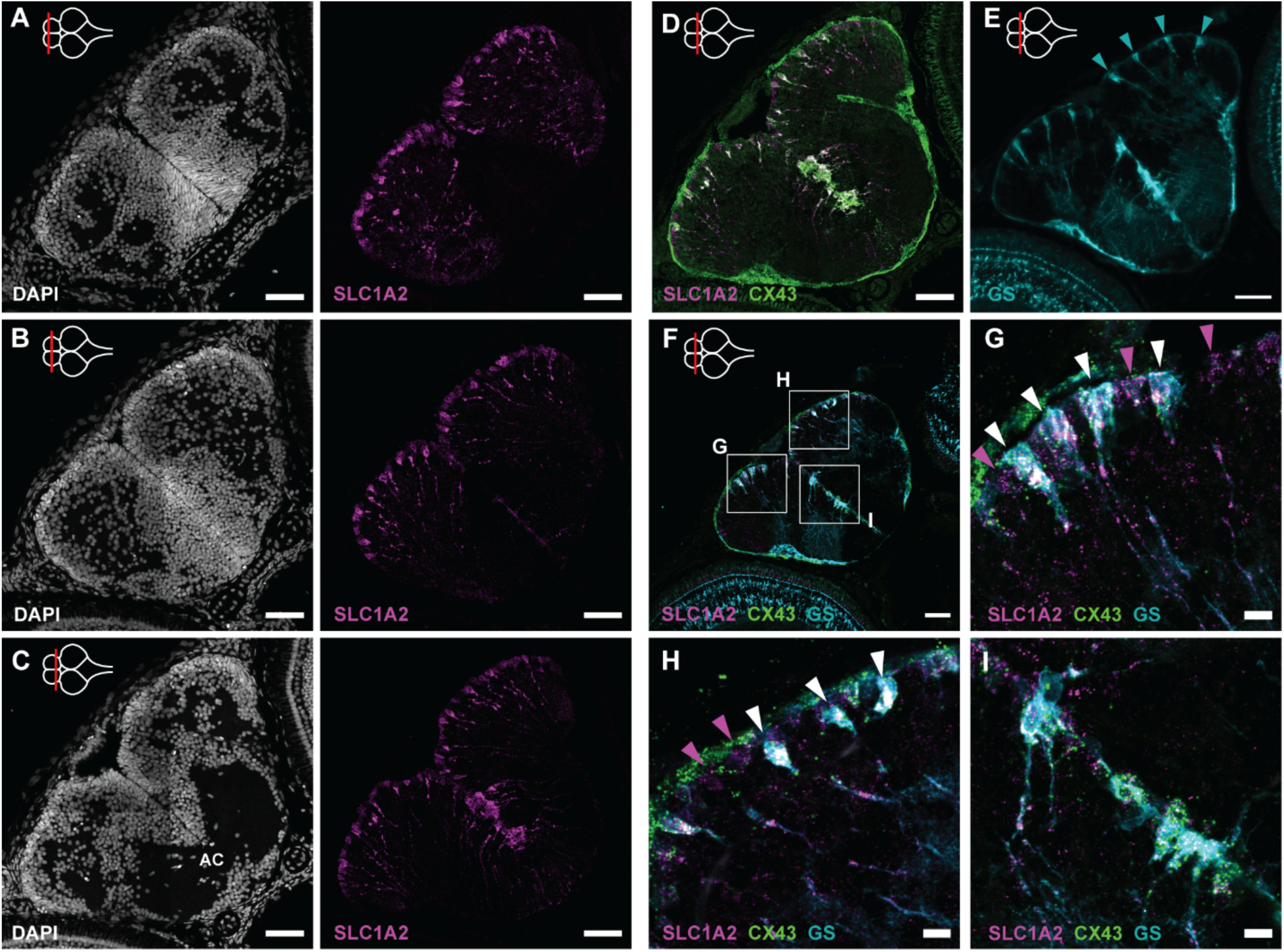
Radial glia patterns in the developing telencephalon. ***(A-C)** Coronal sections of the telencephalon at 5dph along the anterior-posterior axis. Left panels show the nuclear stain DAPI (white) to delineate the nuclear composition and density of the telencephalic domains. Right panels display HCR-FISH targeting SLC1A2 (magenta), an Astro RG1 marker gene in the young-adult telencephalon. Cell bodies, at the ventricular surface, and fiber structures, running towards the pia, are visible on all levels. **(D)** HCR-FISH targeting SLC1A2 (magenta) and CX43 (green), a general Astro-RG marker, on a coronal section of the posterior telencephalon at 5dph. At the midline, a dense cluster of CX43^+^SLC1A2^+^ RGs could be identified as the astroglia subcluster Astro-RG2. **(E)** Immunohistochemical staining of GS (turquoise) on a 5dph telencephalon section at a posterior level. Isolated GS^+^ RGs are positioned one by one at the pallial surface (turquoise arrowheads), and in a dense cluster at the subpallial midline. **(F)** HCR-FISH targeting SLC1A2 (magenta) and CX43 (green) in combination with the immunohistochemical GS (turquoise) staining on a coronal section at the mid-anterior-posterior level. The anterior-posterior position of the sections (A-F) is indicated with a red line on a top view illustration of the brain in the upper left corner of the panels. **(G-I)** Magnification of the squares in panel F. GS^+^/SLC1A2^+^/CX43^+^ RGs (white arrowhead) are interspersed with SLC1A2^+^-only RGs (magenta arrowheads), that are most probably not fully mature yet, at the pallial surface. **(A-F)** Scale bar: 50 µm; **(G-I)*** Scale bar: 10 µm. ***Abbreviation:*** *AC: anterior commissure, HCR-FISH: Hybridization Chain Reaction – Fluorescent In-Situ Hybridisation*.

One of the critical functions of astrocytes in neuronal signaling is the uptake of glutamate by GLT1 at the tripartite synapse (Tanaka *et al*., 1997). Local translation of SLC1A2 mRNA in the RG fibers, visible in all levels of the anterior-posterior axis (Fig. 3A-C), suggests that killifish astroglia-like RGs (Astro-RG1, Astro-RG2) have a similar glutamate-clearing function (Ayana *et al*., 2023). In the killifish posterior subpallium, at the level of the anterior commissure, we could observe a dense cluster of SLC1A2^+^ RGs (Fig. 3C). This cluster of RGs also showed a high expression of CX43 (Fig. 3D). The location and expression pattern of these CX43^+^/SLC1A2^+^ RGs identifies them as the subpallial Astro-RG2 population, already present in the 5dph killifish telencephalon. The fibers of Astro-RG2 followed a specific curvature into the lateral and ventral subpallium (Fig. 3C-D) and strongly resembled a subtype of adult zebrafish RG progenitors with a more oval cell body and thick unbranched processes described before (März *et al*., 2010).

Previous killifish studies used glutamine synthetase (GS) to mark the RGs in the telencephalon (Coolen *et al*., 2020; Van houcke *et al*., 2021). To further delineate the RG subtypes identified by Ayana et al (2023) and compare with previous findings, we combined the expression study of GS with the RG1/RG2 discriminating markers SLC1A2 and CX43. At 5dph, isolated GS^+^ cell bodies appeared positioned one by one at the pallial surface with radial fibers extending to the pial surface of the dorsal pallium (Fig. 3E). Triple-labeling of GS with SLC1A2 and CX43 showed that all GS^+^ cells were positive for both Astro-RG markers (Fig. 3F-I). In between such triple positive RGs (Fig. 3G-H, white arrows), we could observe single SLC1A2^+^ cell bodies (magenta arrows). These may represent an immature or intermediate Astro-RG cell state. At 2w, such GS^+^/SLC1A2^+^/CX43^+^ cell bodies appeared in a more dense group at the pallial surface. Yet, some immature SLC1A2^+^, CX43^-^/GS^-^ cells were present in between the clusters of mature Astro-RGs at the pallial surface (Suppl. Fig. 3B-C). At the subpallial midline, we found a more dense cluster of mature GS^+^/SLC1A2^+^/CX43^+^ RGs already from 5 dph, presumably the Astro-RG2 cluster. In summary, a tilled RG-fiber pattern is visible from hatching in the telencephalon. At the subpallial midline, a dense and mature astroglial cell cluster is present from early development, while a mixture of mature and immature Astro-RG1s line the pallial surface.

### Heterogeneity in the proliferative progenitor populations upon maturation

Next to the Astro-RG subpopulations, a ZIC2^+^ neuroepithelial RG subtype (NE-RG) was identified as the putative root stem cell cluster in the young-adult killifish telencephalon (Ayana *et al*., 2023). NE-RGs are located in subpallial neurogenic region I and pallial neurogenic region III (Ayana *et al*., 2023). Here, we confirmed that ZIC2^+^ NE-RGs were spatially restricted to these neurogenic regions at the ventral subpallial midline and at the posterior part of the lateral pallium from the start of post-embryonic development (Fig. 4A-C, Suppl. Fig. 3D-G). In both regions, the NE-RGs expressed typical progenitor markers at the ventricular border (ZIC2^+^, SOX2^+^). Further into the parenchyme, these cells lost SOX2 expression in the pallium (ZIC2^+^, SOX2^-^) but not in the subpallium. (Fig. 4D-E). At the pallial-subpallial border, a dense cluster of SOX2^+^ postmitotic cells was present in the parenchyme (Fig. 4B-C).

**Figure 4:**
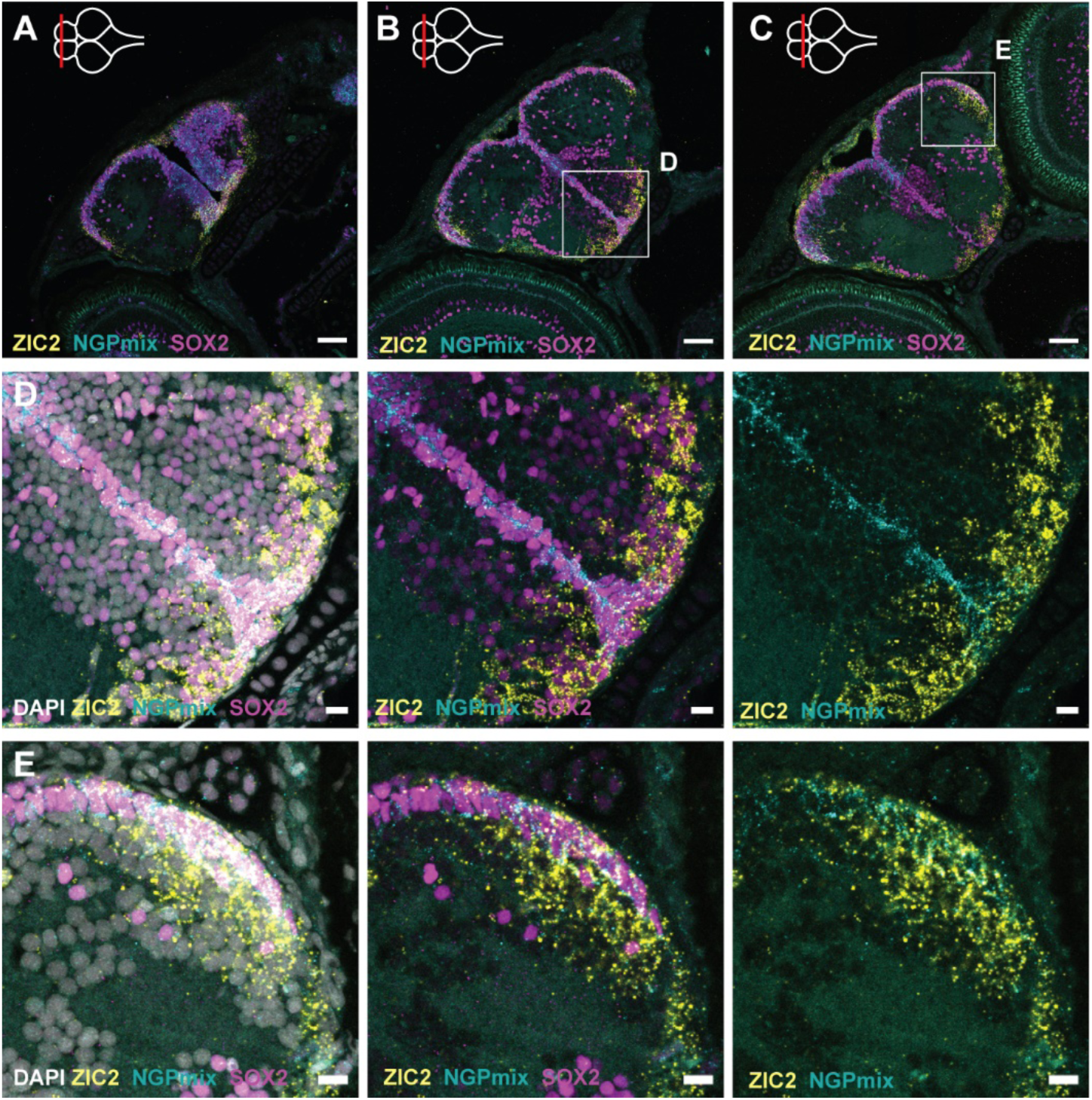
Distribution of the non-glial and neuroepithelial-like progenitors. **(A-C)** HCR-FISH targeting NGPmix (=STMN1A + HMGB2A, NGPs, turquoise) and ZIC2 (NE-RGs, yellow) in combination with an immunohistochemical staining for the progenitor marker SOX2 (magenta) on coronal sections along the anterior-posterior axis in the 5dph telencephalon. Scale bar: 50 µm. **(D-E)** Magnifications of the squares in B and C. The HCR-FISH targeting ZIC2 (yellow) and NGPmix (turquoise) and the SOX2 staining (magenta) are combined with the nuclear stain DAPI (white). NGPs and NE-RGs are present in the subpallial midline and posterior pallial neurogenic regions. In the subpallium, both cell types retain their progenitor profile (SOX2^+^), while in the pallium, NE-RG3 loses its progenitor profile (SOX2^-^) in the parenchyme. Scale bar: 10 µm.

In the killifish telencephalon, it are the NGPs, and not the RG populations, that are responsible for the bulk of proliferation from post-embryonic development until adulthood (Coolen *et al*., 2020; Van houcke *et al*., 2021; Ayana *et al*., 2023). In the developing post-embryonic killifish, we could identify the NGPs close to NE-RGs up until 2w (Fig. 4, Suppl. Fig. 3D-G). This strengthens the idea of a lineage relationship during development between these progenitor types and confirms the trajectory analysis in the young-adult killifish telencephalon (Ayana *et al*., 2023).

To characterize the proliferative profile of all major progenitor subtypes in the developing killifish telencephalon, we examined the spatial distribution of the proliferation marker PCNA (Fig. 5, Suppl. Fig. 4). Previously, Coolen *et al*. (2020) already investigated the proliferation capacity of GS^+^ RGs and SOX2^+^ GS^-^ NGPs in the developing killifish pallium. They identified a remarkable reduction of dividing GS^+^ RGs in the first 2w of post-embryonic development. We probed for double labeling of PCNA with the Astro-RG, NE-RG and NGP discriminating markers CX43, ZIC2, and the combination of STMN1A and HMGB2A (referred to as NGPmix) in the 1dph, 5dph and 2w telencephalic pallium and subpallium (Fig. 5A-B, Suppl. Fig. 4A, C). At 5dph, we could observe a general co-labeling between NGPmix and PCNA, validating the highly proliferative nature of NGPs in the developing telencephalon. In line, the NGPs were proliferative from 1dph until 2w in all three neurogenic regions (Fig. 5C-E, Suppl. Fig. 4A-C). The NE-RGs at the pallial ventricular border were proliferative (ZIC2^+^, PCNA^+^), but the NE-RGs extending into the parenchyme did not express PCNA, in line with the absence of SOX2 expression in these cells (Fig. 4E, ig. 5E, Suppl. Fig. 4D). In the subpallium, we observed both PCNA^+^ and PCNA^-^ NE-RGs. Of note, the NE-RGs in the subpallium lost their proliferative capacity (PCNA^-^) but kept SOX2 expression outside of the proliferative region (Fig. 4D, Fig. 5C). Comparable patterns of dividing and non-dividing (quiescent) NE-RGs were found during the first two weeks of development and are also comparable to previous findings in the young-adult telencephalon (Ayana *et al*., 2023).

**Figure 5:**
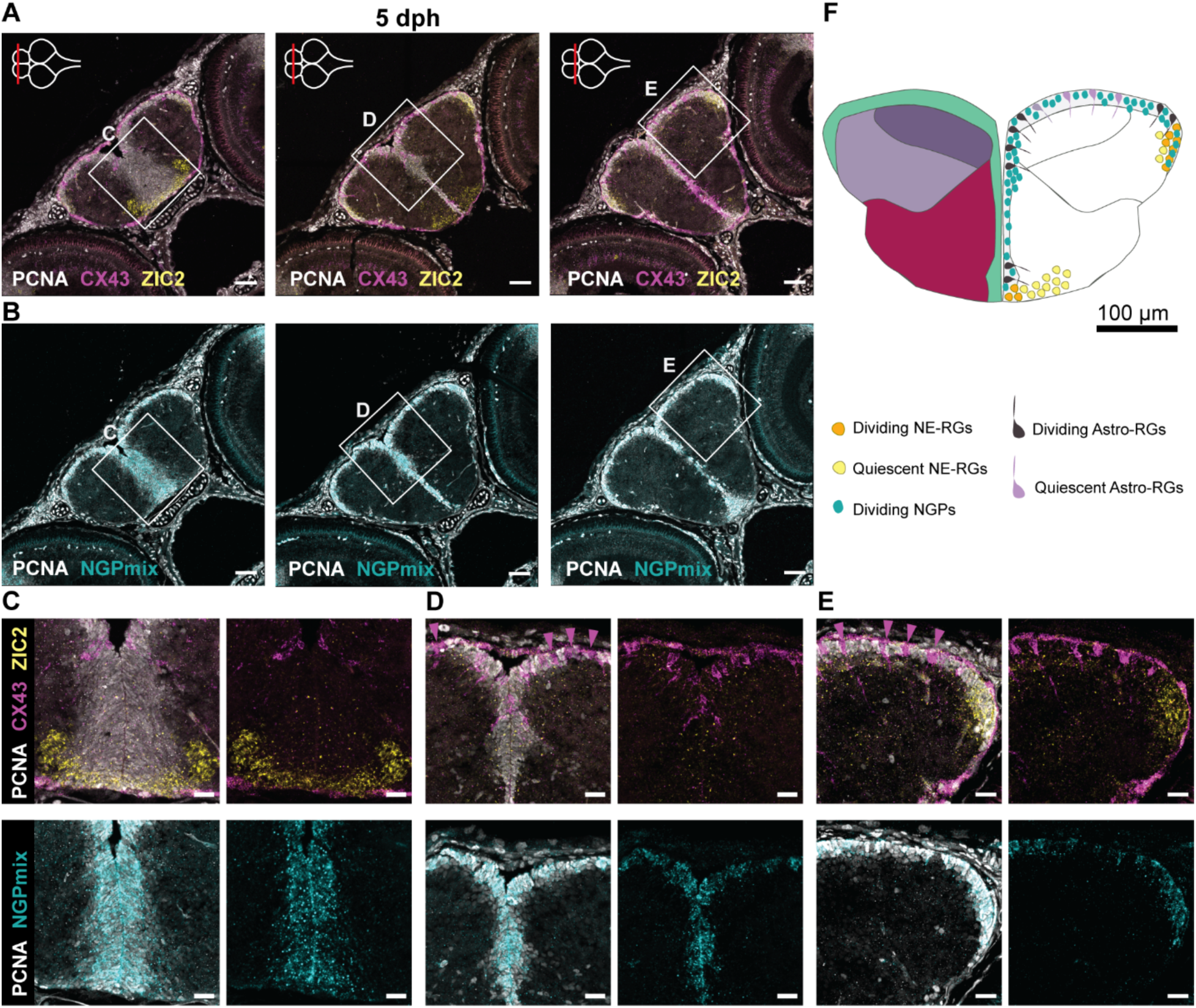
Dynamic progenitor heterogeneity in the developing telencephalon. **(A-B)** Adjacent coronal sections of the telencephalon at 5dph along the anterior-posterior axis. The level of each section is indicated with a red line on the top view brain illustration in the upper left corner of each panel in A. HCR-FISH targeting (A) CX43 (pan Astro-RG, magenta) and ZIC2 (NE-RG, yellow) or (B) NGPmix (turquoise), is combined with an immunohistochemical staining for the proliferation marker PCNA (white). Scale bar: 50 µm. NGPs account for the bulk of proliferation, Astro-RGs and NE-RGs appear both dividing and non-dividing depending on the location in the telencephalon **(C-E)** Magnification of the squares in A-B. Non-dividing Astro-RGs (CX43^+^, PCNA^-^) are indicated with a magenta arrowhead and appear isolated in between stretches of dividing NGPs. Scale bar: 20 µm. **(F)** Illustration of a 5dph coronal telencephalic section at mid-anterior-posterior level. The distribution of dividing and non-dividing Astro-RGs, NE-RGs and NGPs is displayed in the ventricular zone.

The role of Astro-RGs in early brain growth (5dph) seemed to be confined to one type of proliferating Astro-RGs (PCNA^+^, CX43^+^) that were mainly located at the dorsal midline and close to the posterior pallium, while non-proliferating Astro-RGs (PCNA^-^, CX43^+^) were located at the dorsal pallial surface (Fig. 5D-E). The posterior subpallial Astro-RG cluster, previously identified as Astro-RG2, also showed no proliferation (Fig. 5A). At 2w, a strong decrease in proliferating Astro-RGs was observed, also at the dorsal pallial midline (Suppl. Fig. 4C,E). In summary, the killifish progenitor pool exists out of the highly proliferative NGPs, robust populations of dividing and quiescent NE-RGs in specific niches, a mature subpallial Astro-RG cluster and a pallial Astro-RG progenitor population with a quick decline in proliferation after hatching (Fig. 5F).

### Intermediate progenitors in the pallial ventricular region

Intermediate progenitors are RG daughter cells in the embryonic mouse cerebral cortex and are an important secondary proliferating progenitor population generating the bulk of excitatory glutamatergic projection neurons. These intermediate progenitors specifically express EOMESA (TBR2) (Hevner, 2019). In the adult zebrafish telencephalon, expression of EOMESA is observed in most parenchymal neuronal zones and at the pallial ventricular layer (Ganz *et al*., 2015). Here, we explored the presence of EOMESA^+^ progenitors in the neurogenic niches of the developing killifish telencephalon. EOMESA^+^ cells could be found at the ventricular border of the pallial surface (Fig. 6A-D). To investigate whether these cells were direct descendants of NGP or RG cells, we reasoned that we would find marker co-expression in a portion of these cells if they shared the lineage. Double-labeling with the NGP markers (STMN1A and HMGB2A) revealed co-expression of EOMESA and NGPmix in the neurogenic niches (Fig. 6B-D). This was particularly visible at the pallial midline in the telencephalon at 2w, where EOMESA expression did not extend into the parenchyme but was restricted to these cells at the ventricular border (Suppl. Fig. 5A-C). We additionally probed for co-expression of the Astro-RG marker CX43 and EOMESA but could not detect any. Still, EOMESA^+^ cells were found in close proximity to CX43^+^ Astro-RGs. This was very pronounced in the 2w telencephalon, where there is a higher abundance of mature Astro-RGs (Fig. 6B-D, Suppl. Fig. 5B-C). So, EOMESA^+^/NGPmix^+^ intermediate progenitors could be found in the pallial ventricular regions, intermingled with NGPs (NGPmix^+^) and Astro-RGs (CX43^+^). In the anterior subpallium, we could also observe EOMESA^+^ cells at the outer border of the ventral NGP-rich neurogenic region (Fig. 6D, Suppl. Fig 5A). Since intermediate progenitors have the capacity to divide a limited number of times before differentiating into glutamatergic neurons, we verified the proliferation profile of these EOMESA^+^ cells in the neurogenic regions. Double-labeling with the proliferation marker PCNA, confirmed that the EOMESA^+^ cells in the progenitor regions of the telencephalon can proliferate at hatchling and juvenile stages (Fig. 6E-F, Suppl. Fig. 5D-E). Our results suggest that the EOMESA^+^ intermediate progenitors in the developing killifish telencephalon presumably arise from NGPs and less likely from RGs, contrary to what is observed in mammals and zebrafish (Furlan *et al*., 2017; Hevner, 2019; Labusch *et al*., 2020). Accordingly, an NGP subpopulation in the dorsal pallium seems to be committed to the glutamatergic neurogenic lineage.

**Figure 6:**
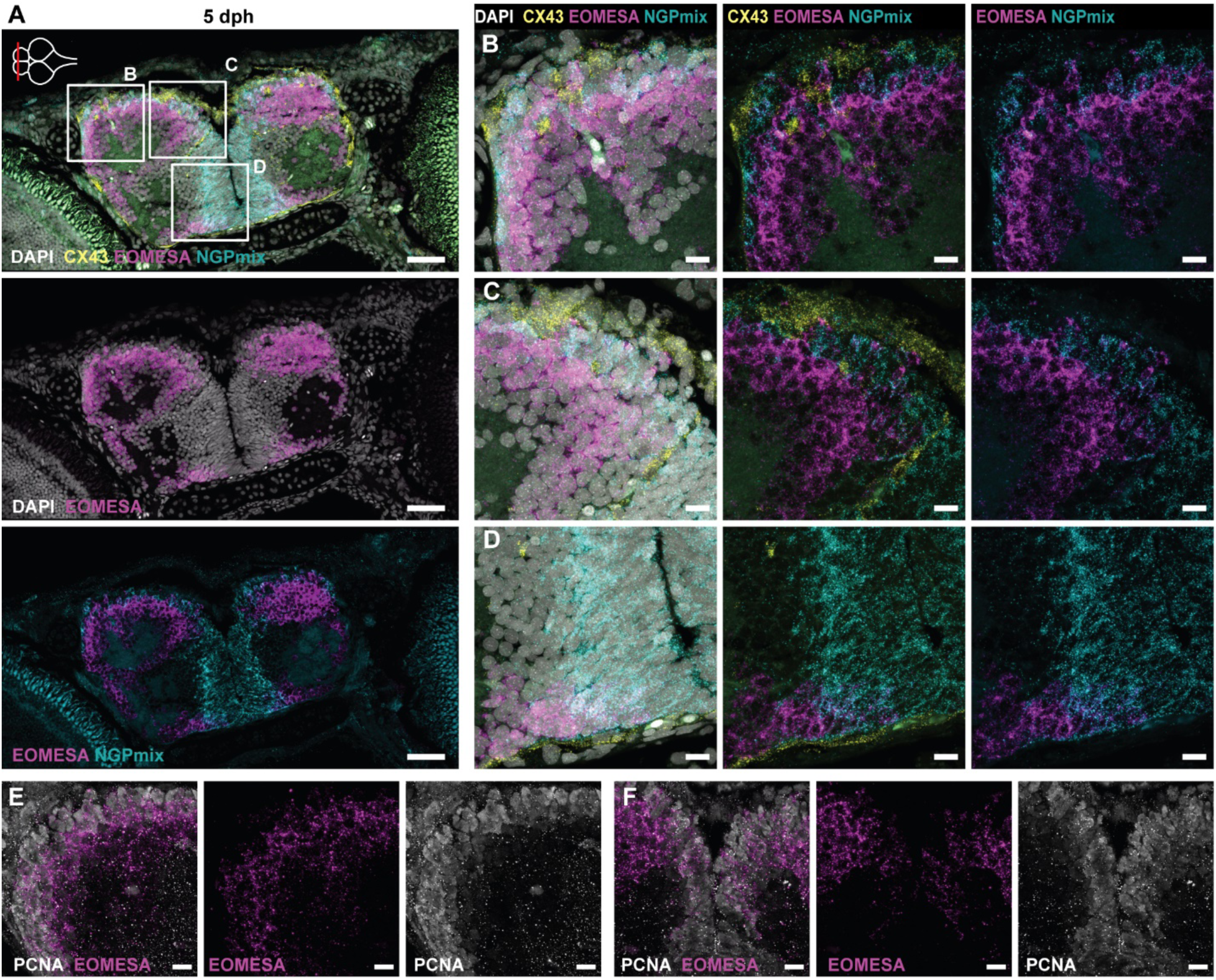
Intermediate progenitors in the killifish pallium. **(A)** HCR-FISH targeting CX43 (Astro-RGs, yellow), NGPmix (turquoise) and the immature excitatory neuron marker EOMESA (magenta) in combination with the nuclear stain DAPI (white) on a coronal section of the telencephalon at 5dph. The anterior-posterior level is depicted with a red line on the top view brain illustration in the upper left corner. Scale bar: 50 µm. **(B-D)** Magnification of the squares in A. Intermediate progenitors (EOMESA^+^/NGPmix^+^) are found at the pallial neurogenic region and ventral subpallium, dispersed in between Astro-RGs (only in pallium) and NGPs. Scale bar: 10 µm. **(E-F)** HCR-FISH targeting EOMESA (magenta) in combination with an immunohistochemical staining for the proliferation marker PCNA (white). The panels are magnifications of zones on a section of a comparable anterior-posterior level as in A. E and F are magnifications of the lateral pallium and dorsal midline, respectively. Proliferating PCNA^+^/EOMESA^+^ progenitors are present at the midline and pallial surface. Scale bar: 10 µm.

### Development and organization of inhibitory and excitatory neurons

To visualize where neuronal subtypes are born in the post-embryonic killifish, we probed for the presence of two main neuronal classes: gamma-aminobutyric acid-releasing (GABAergic) neurons and glutamatergic neurons, using the canonical markers GAD1B and SLC17A7 (VGLUT1), respectively. To examine the maturation state of these post-mitotic cells, we combined this with the immature inhibitory and excitatory neuron markers DLX1 and EOMESA respectively. DLX1 is known to be an important transcription factor for neuronal differentiation and migration of GABAergic neurons (Cobos *et al*., 2007).

DLX1^+^ cells arise next to the NGP-rich neurogenic region in the subpallium (Fig. 7A-C). DLX1^+^/GAD1B^+^ and DLX1^-^/GAD1B^+^ inhibitory neurons were heavily populating the ventral portion of the rostral telencephalon (Fig. 7A, G). GAD1B^+^ cells could also be found scattered throughout the pallium. The absence of DLX1^+^ cells in the pallial neurogenic niches indicated that these GAD1B^+^ GABAergic neurons were formed solely in the subpallium and likely arrived in the pallium via tangential migration from the subpallium. During zebrafish development, invasion of GABAergic neurons in the pallium is observed from 2-3 dpf, however, the scattered pattern of GAD1B^+^ cells in all subdivisions of the pallium as observed here in the killifish, could only be visualized in zebrafish with GABA from 3 dpf (Mueller *et al*., 2006, 2008). In the mid-posterior telencephalon (Figure 7B), a comparable pattern of DLX1^+^ and GAD1B^+^ neurons could be observed, albeit at a lower density. Of note, the posterior subpallium showed a lower density of cell bodies in general (Fig. 1B). GAD1B^+^ cells were also found at the lateral border of the mid-posterior subpallium. At the posterior edge of the telencephalon, the ventral subpallium (preoptic area) showed a high density of DLX1^+^ and GAD1B^+^ cells at the midline (Fig. 7C).

**Figure 7:**
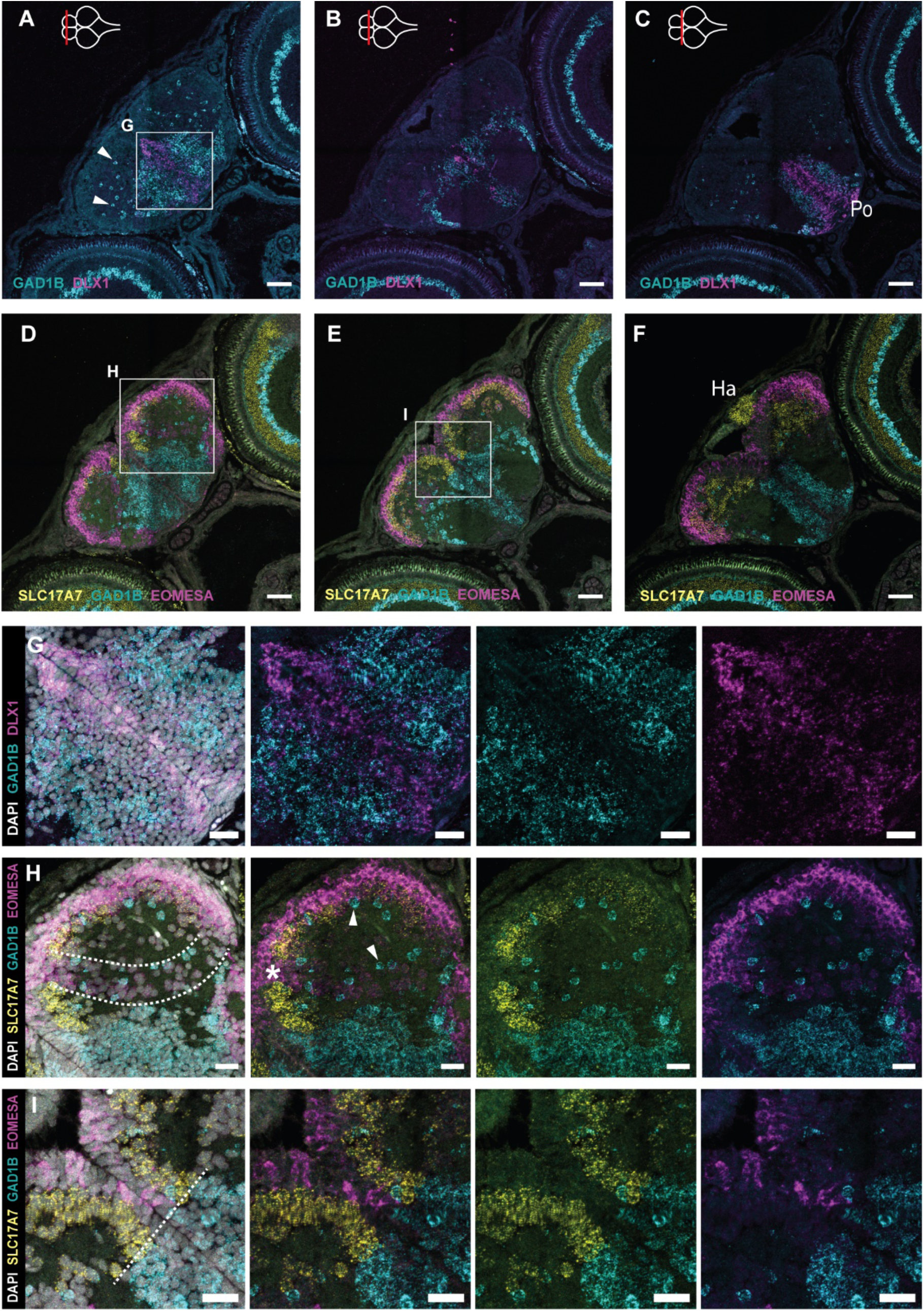
Excitatory and inhibitory neuron clusters in the 5dph telencephalon. **(A-C)** Coronal sections of the telencephalon at 5dph along the anterior-posterior axis. HCR-FISH targeting DLX1 (magenta) and GAD1B (turquoise) mRNA expression shows the distribution of immature and mature inhibitory neurons, respectively. **(D-F)** Adjacent coronal sections of A-C, HCR-FISH targeting EOMESA (magenta) SLC17A7 (yellow) and GAD1B (turquoise) mRNA expression shows the distribution of immature and mature excitatory neurons, respectively, and the mature inhibitory neurons. The anterior-posterior position of the sections is indicated with a red line on the top view brain illustration in the upper left corner of panels A-C. The border of the telencephalon (posterior-most section C,F) is recognizable based on the presence of the habenula (Ha) on top of the dorsal pallial surface, positive for SLC17A7. **(G)** Magnification of the square depicted in panel A. HCR-FISH labeling DLX1 and GAD1B in combination with a nuclear stain (DAPI, white) to visualize all cell bodies. Progenitor cells at the midline are negative for DLX1 (DAPI^+^). **(H-I)** Magnification of the squares depicted in panels D-E. HCR-FISH labeling EOMESA, SLC17A7 and GAD1B in combination with DAPI. The progenitor cells at the ventricular surface are negative for EOMESA (DAPI^+^). **(H)** A stream of low-level EOMESA^+^ cells is visible (area between dotted lines), stretching from medial to ventrolateral pallium. This band is devoid of SLC17A7^+^ expression (asterisk). **(I)** A clear border is discernable between inhibitory (GAD1B^+^) and excitatory (SLC17A7^+^ and/or EOMESA^+^) cells around the midline, indicated with a dotted line. GAD1B^+^ cells are found scattered throughout the pallium (white arrowheads). **(A-F)** Scale bar: 50 µm **(G-I)** Scale bar: 20 µm. **Abbreviations:** Ha: Habenula, Po: Preoptic area.

EOMESA^+^ neurons could be observed at the periventricular zone of the pallium along the anterior-posterior axis (Fig. 7D-F, Suppl. Fig. 6A-C). More mature excitatory neurons (SLC17A7^+^) were positioned at the parenchymal side of the immature EOMESA^+^ neurons, in line with the stacking growth model of the pallium. The neural progenitor cells at the midline and ventricular surface, lacking neuronal markers, can be clearly distinguished from the cells committed to the neuronal lineage (Fig. 7G-H). Anteriorly, we could observe a stream of EOMESA^+^ neurons extending from the medial pallium to the posterior pallial region (Fig. 7H, in between dotted lines). This stream was devoid of mature excitatory neurons (Fig. 7H, asterisk), but GAD1B^+^ neurons were scattered throughout. GAD1B^+^ mature inhibitory neurons could also be found near SLC17A7^+^ mature excitatory neurons from 1dph onwards (Suppl. Fig. 6D, Fig. 7H). The pallial-subpallial border could be distinguished by the clear expression boundary between inhibitory and excitatory cells at the midline (Fig. 7I). In sum, the early post-hatching killifish has a patterned telencephalon with glutamatergic neurons generated at the pallial surface and GABAergic cells in the subpallium.

## Discussion

The African turquoise killifish is an extensively used model for aging research in a broad spectrum of research fields, including genetics, evolution, immunology and neurobiology. We and others have exploited its conveniently short lifespan to study the aging process of the central nervous system, emphasizing the impact on neuro(re)genesis. Here, we probed when the progenitor population sustaining adult neurogenesis in the killifish telencephalon arises, and how the explosive growth process during early post-hatching development takes place. We charted the sequential generation of neuronal cells and telencephalon morphogenesis from hatchling to young-adult stages and identified: (1) a distinct stacking process of newborn cells in the pallium and subpallium, (2) a more lateral and faster growth of the pallium than the subpallium post-hatching, (3) the presence of a maturing scaffold of RG nature early post-hatching, (4) a high proliferative activity of progenitor cells, and a cell type and region dependent decrease in proliferation over time, and (5) a clear pallial-subpallial patterning of progenitor cells committed to either the glutamatergic or GABAergic neuronal lineage. The spatial map presented here shows the diversity of progenitor cells, constructed during development and maintained in adulthood, and will be indispensable in future studies on lineage relationships, and age-related decline in neurogenesis and regeneration.

A study on whole body growth of GRZ killifish reported accelerated growth in the first four weeks after hatching with a peak in the second week (Blažek *et al*., 2013). The growth layers in the pallium and subpallium, visualized with EdU birthdating, revealed a comparable growth curve for the telencephalon specifically. The growth from 4w onward is negligible compared to the thickness of the layers produced earlier in development. Measurements of the dorsoventral and mediolateral growth of the telencephalon also revealed that from 2w to young-adult (6w), the pallium still doubled in size on average in all directions measured. For the subpallium, this difference was smaller.

The EdU tracing experiment revealed a distinct stacking process for the pallium and subpallium. Comparable experiments in other teleost models mainly focused on the pallial growth. In zebrafish, a similar stacking process of the pallium from embryonic to juvenile stages was revealed, using a Tet-On birth dating strategy to specifically trace the progeny of HER4^+^ RGs (Furlan *et al*., 2017). In teleost fishes that have an inverted telencephalon, the central pallial core, created during embryogenesis, thus becomes surrounded by concentric layers of newborn cells generated sequentially over time. In zebrafish, HER4^+^

RGs produce the majority of newborn neurons (Furlan *et al*., 2017). In the developing zebrafish, however, two neural progenitor lineages are identified in the pallium. One emerges from HER4^+^ RGs at the dorsomedial pallium, giving rise to neurons and new RGs. The other originates at the lateral pallium and has a neuroepithelial and amplifying character which thus resembles the NE-RG progenitors discovered in killifish (Dirian *et al*., 2014; Ayana *et al*., 2023). Than-Trong and colleagues (2020) also described a HER4^-^ progenitor pool as the source of the gradual expansion of the RG population in the adult zebrafish (Than-Trong *et al*., 2020). While HER4^+^ (HES5) RGs are also found in killifish telencephalon, expansive growth is supported by NGPs in all neurogenic regions during post-embryonic development and in the adult killifish telencephalon (Coolen *et al*., 2020; Ayana *et al*., 2023). The spatial proximity of NGP and NE-RG marker expression at the subpallial midline and posterior pallium is prominent around hatching (1-5dph), suggestive of a lineage relationship between these progenitor subtypes from the early stages onwards. Lineage tracing from NE-RG3 and NGPs is necessary to identify the exact progeny of these cell types and renewal of the progenitor pool during development, adulthood and aging.

A scaffold-like structure of radial fibers is already present from hatching in the killifish telencephalon. The non-overlaying radial structure is reminiscent of tiling of RGs and astrocytes in the developing and mature mouse brain (Bushong *et al*., 2002; Nakagawa *et al*., 2019). Tiling of glial cells, well described in mammals for cortical astrocytes, is a conserved mechanism already present in invertebrates (Baldwin *et al*., 2021; Pogodalla *et al*., 2022). Although present from 1dph, the radial scaffold might still be maturing upon hatching, as immature GS^-^ Astro-RG1s were found interspersed between mature Astro-RG1s. In the subpallium, on the other hand, a dense cluster of mature subpallial Astro-RG2s is populating the midline already at hatching. These Astro-RG2s are quiescent and express the mature astrocyte markers CX43, SLC1A2 and GS at 1dph. Future studies are necessary to reveal if these early astrocyte-like cells originate directly from progenitor RGs, as described in mammals, or via the NGPs, the predominant progenitor type in the post-embryonic killifish subpallium. M. Coolen *et al* (2020) identified a strong decrease in proliferating RGs at the dorsal surface of the killifish pallium. Our data revealed that these RGs represent the pallial Astro-RG type, Astro-RG1. We validated the decrease in proliferating Astro-RG1s and discovered an uneven distribution of dividing and quiescent pallial Astro-RG1s. The dividing Astro-RG1s seem to be confined to the dorsal midline and posterior pallium, but their number is relatively limited and the bulk of proliferation comes from the NGPs.

Using deterministic markers (DLX1, EOMESA, GAD1B, SLC17A7) we could visualize the maturation sequence of excitatory and inhibitory neurons in the killfish telencephalon. The subpallium is already densely populated with mature GABAergic neurons at hatching. At 1dph, mature GABAergic neurons are also integrated into the pallium, suggestive of active migration from the subpallium to the pallium. Neurons from the GABAergic lineage are exclusively generated in subpallial neurogenic Region I, homologous to the mammalian subventricular zone. Progenitors and immature neurons committed to the glutamatergic lineage were identified in both the pallial and subpallial neurogenic regions. Subpallial EOMESA^+^ cells were spatially restricted to the anterior-ventral most part of the telencephalon. In other teleost species, the ventral and lateral nuclei of the anterior subpallium are thought to be homologous to the septum and known to have expression of EOMESA (Wullimann and Mueller, 2004; Mueller *et al*., 2008). In addition to TBR^+^ (EOMESA) intermediate progenitors in the developing mouse neocortex, TBR2^+^ progenitors were identified, involved in the production of glutamatergic neurons for the lateral olfactory tract (Englund *et al*., 2005; Vasistha *et al*., 2015). Vasistha *et al* (2015) also confirmed that TBR2^+^ progenitor cells exclusively produce glutamatergic neurons and not GABAergic neurons or astrocytes. In the killifish pallium, glutamatergic neurons are produced along the entire dorsal surface. We indirectly identified radially migrating neurons, creating the layered inside-out pallial structure, and laterally migrating neurons, comparable to what is described in the developing zebrafish (Mueller *et al*., 2008). Even though the timing of both embryonic and post-embryonic development is considerably different for zebrafish and killifish, we observed a comparable spatial organization of inhibitory and excitatory neurons for the telencephalon once hatched (Wullimann and Mueller, 2004; Mueller *et al*., 2006, 2008, 2011; Ganz *et al*., 2015). Our study reveals the neuronal diversity in the developing killifish telencephalon by spatially delineating the immature and mature excitatory and inhibitory neurons. In the young-adult killifish telencephalon, single-cell sequencing revealed a more extensive diversity of cell types, including 12 excitatory and 5 inhibitory neuronal clusters (Ayana *et al*., 2023). The advent of large-scale single-cell studies is likely to discover an even more impressive diversity, such as the recently generated neuronal cell type atlas of the goldfish telencephalon, generated by combining scRNA-sequencing and spatial transcriptomics, which uncovered 88 GABAergic and glutamatergic neuronal subclusters (Tibi *et al*., 2023). In conclusion, we revealed the cellular diversity during explosive growth of the killifish telencephalon, where we uncovered specific progenitor signatures for each neurogenic region and evidence for distinct glutamatergic and GABAergic progenitor lineages in the post-embryonic telencephalon.

## Acknowledgment

We thank Simon Buys and Arnold Van Den Eynde, and the KU Leuven killifish team in general, for taking care of the fish facility. We want to thank Lieve Geenen for her substantial help during tissue processing. We also acknowledge the invaluable help of Ruth Styfhals in introducing the Hybridization Chain Reaction technique in the lab.

## Author contributions

**C.Z**.: Conceptualization, Methodology, Investigation, Formal analysis, Writing: original draft preparation, review and editing, Visualization. **V.M., A.R., J.V.:** Writing: review and editing. **L.A., E.S.**: Supervision, conceptualization, Funding acquisition, writing: review and editing.

